# AUC-PR is a More Informative Metric for Assessing the Biological Relevance of In Silico Cellular Perturbation Prediction Models

**DOI:** 10.1101/2025.03.06.641935

**Authors:** Hongxu Zhu, Amir Asiaee, Leila Azinfar, Jun Li, Han Liang, Ehsan Irajizad, Kim-Anh Do, James P. Long

## Abstract

In silico perturbation models, computational methods which can predict cellular responses to perturbations, present an opportunity to reduce the need for costly and time-intensive in vitro experiments. Many recently proposed models predict high-dimensional cellular responses, such as gene or protein expression to perturbations such as gene knockout or drugs. However, evaluating in silico performance has largely relied on metrics such as *R*^2^, which assess overall prediction accuracy but fail to capture biologically significant outcomes like the identification of differentially expressed genes. In this study, we present a novel evaluation framework that introduces the AUC-PR metric to assess the precision and recall of DE gene predictions. By applying this framework to both single-cell and pseudo-bulked datasets, we systematically benchmark simple and advanced computational models. Our results highlight a significant discrepancy between *R*^2^ and AUC-PR, with models achieving high *R*^2^ values but struggling to identify Differentially expressed genes accurately, as reflected in their low AUC-PR values. This finding underscores the limitations of traditional evaluation metrics and the importance of biologically relevant assessments. Our framework provides a more comprehensive understanding of model capabilities, advancing the application of computational approaches in cellular perturbation research.

## Introduction

Cellular perturbation experiments play a fundamental role in modern biological and medical research. In these experiments, the normal state of living cells is deliberately altered through various perturbations. These perturbations can be classified into several categories, including genetic (gene knockdown, gene knockout, gene overexpression), chemical, physical[11], and metabolic[30]. By comparing perturbed and unperturbed cellular states, researchers can study perturbation effects [1, 25, 31] and infer the function of targeted genes, proteins, and other cellular components [21, 24].

Various methods are employed to assess cellular responses to perturbations, including gene expression analysis [12, 15], protein activity measurements [6, 23], and cellular morphology studies [20]. Among these response types, gene expression profiling has emerged as the predominant measurement approach due to its extensive downstream biological applications [26]. The results from these experiments have wide-ranging applications in drug discovery and development. Specifically, perturbing genes or proteins helps identify their roles in diseases, thereby guiding the development of targeted therapies [25]. Furthermore, by manipulating gene expression, scientists gain crucial insights into gene functions within cellular pathways, advancing our understanding of biological systems and therapeutic strategy development. These approaches have proven particularly valuable in identifying combination therapies for diseases such as cancer [33], where understanding drug synergies and antagonistic effects can lead to more effective treatment strategies.

However, the high costs associated with exhaustive experimental studies make it impractical to explore every possible outcome in factorial experiments where all perturbations are applied to all cellular variations in vitro. For instance, the LINCS program [15] encompasses data from 71 cell lines and over 25,000 perturbations, including small molecule compounds, gene knockdowns or overexpressions, and biologics. Despite this extensive scope, fewer than 10% of the approximately 1.75 million potential experiments were conducted, highlighting the substantial resource demands of such large-scale studies.

To address this challenge, in silico models—computational methods designed to predict cellular responses to untested perturbations based on historical experimental data—have been developed. By leveraging these predictive models, researchers can estimate cellular behaviors without the need for physical experiments. This approach simplifies the research process and significantly reduces the costs associated with conducting large-scale experimental studies. Several studies, such as [17], [18] and [19] introduced variational autoencoder-based methods to predict out-of-sample single-cell perturbation responses. Building on this, [29] enhanced these predictions by integrating prior knowledge with multi-layer neural networks to forecast post-perturbation scRNA-seq gene expressions. [32], [22] and [28] approach the prediction task from a causal modeling perspective, which aims to infer cause-effect relationships rather than relying purely on statistical associations. Most recently, works that incorporate transformer based large language models like scGPT [8] leverage generative pre-training to learn biological embeddings for predicting genetic perturbation responses, annotating cell types, and integrating multi-omic datasets.

Parameters in these models are learned on a set of perturbations tested in vitro. In most of these works, model performance has been assessed by comparing predicted cellular responses, like gene expressions to the actual responses under the same conditions. Commonly used metrics include *R*^2^ (squared Pearson’s correlation) and mean squared error, with high correlation and low error often cited as evidence of model quality. This evaluation strategy is popular for assessing high-dimensional continuous outcomes [4, 5]. However, while these metrics are useful, this approach lacks scientific interpretability, providing limited insight into the biological relevance of the predictions. For example, a high *R*^2^ score may indicate that a model captures global trends in gene expression but does not ensure that biologically meaningful changes, such as differentially expressed genes (DEGs), are accurately identified.

In practice, the most scientifically important outcome of perturbation experiments is the identification of signature gene or protein markers that are differentially expressed under perturbed conditions compared to unperturbed ones [15]. These signature markers are further used for analyses such as gene set enrichment analysis (GSEA) to study potential causal relationships with diseases. Statistically, differentially expressed markers are determined using both p-values and log fold-changes when comparing gene expressions across conditions. Various methods have been proposed for computing p-values, including GLM-based approaches like ZINB, ZITweedie, GLMM-based methods, along with simpler methods such as t-tests and non-parametric alternatives [9].

For predictions generated by in silico models, a more interpretable evaluation of model performance would be to assess whether the models can accurately identify these signature gene markers. To our knowledge, there has been no research comparing the results of an in vitro differential expression analyses with to an in silico differential expression analysis. This represents a significant shortcoming, as the identification of DEGs is a primary objective of these experiments. In particular, models optimized for overall gene expression prediction may fail to capture the subset of genes most relevant to biological hypotheses, limiting their utility in downstream analyses.

In this work we make the following contributions:

- Present the first instance of performing differential expression analysis using in silico predicted perturbation responses from several perturbation prediction models from [19], [32] and benchmark linear models.
- Propose performance measures to quantify the difference between results of in vitro differential expression analysis and results of in silico differential expression analysis.
- Apply our proposed evaluation method to two public datasets, systematically evaluating in silico model performance using the proposed method on scRNA-seq data and psedo-bulked data.

Our work reveals a discrepancy between traditional evaluation metrics like *R*^2^, and our proposed metric. This finding underscores the limitations of the popular *R*^2^ metric and emphasizes the need for evaluation methods that offer a more biologically meaningful assessment of model performance. By focusing on the model’s ability to detect DEGs rather than overall correlation with experimental data, we introduce a framework that aligns more closely with biological applications.

## Materials and methods

### Mathematical Formulation of Cellular Perturbation Experiments

We present a mathematical framework for modeling and predicting cellular responses to perturbations, with particular emphasis on gene expression. Following [32], we define cellular variations as **contexts** (denoted by *c* ∈ 𝒞, e.g., cell types or cell lines) and perturbations as **actions** (denoted by *a* ∈ 𝒜, e.g., gene knockouts or drug treatments). Each context-action pair (*c, a*) defines an experiment. For each context *c*, we observe the baseline unperturbed state, denoted by the special action *a* = 0. Thus, the complete action space is 𝒜 = 𝒜_*p*_ ∪ {0}, where 𝒜_*p*_ represents the set of perturbations and 0 denotes the control condition.

For a given experiment (*c, a*), let ℱ*ca* denote the *p*-dimensional joint distribution function of gene expressions, where *p* represents the number of genes. This distribution captures both biological variability and experimental noise at the single-cell level. We define the true response for experiment (*c, a*) as the expected value of this distribution:

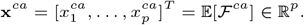

The collection of all true responses can be organized into a third-order tensor:

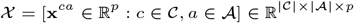

In practice, we observe gene expression data through single-cell RNA sequencing (scRNA-seq) technology [26], which provides cell-level measurements. For experiment (*c, a*), we obtain *n*^*ca*^ independent and identically distributed (i.i.d.) samples (i.e., expression profile of single cells) from ℱ^*ca*^. Let 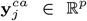 denote the observed expression vector from the *j*-th cell under experiment (*c, a*), where *j* ∈ {1, …, *n*^*ca*^}. We can write:

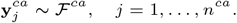

The collection of all cell-level observations for experiment (*c, a*) is denoted by:

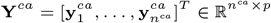

Given these observations, one can estimate the true response **x**^*ca*^ from in vitro experimental data using the sample mean:

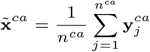

By the Law of Large Numbers, 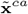 converges to the true response **x**^*ca*^ as *n*^*ca*^ → ∞.

In practice, we can only observe samples from a subset of all possible context-action pairs. Let Ω ⊂ 𝒞 × 𝒜 represent the set of observed experiments, and define the observation set as:

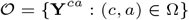

Given these observations, in silico models aim to predict responses for unobserved context-action pairs. We denote such a model as *M* (𝒪, ℛ), where ℛ represents auxiliary information beyond contexts and actions (e.g. chemical structure of drug perturbations). Different models may produce different outputs:

- **Mean Response Prediction:** Some models [32, 22] predict only the mean response **x**^*ca*^ ∈ R^*p*^ for a target pair (*c, a*).
- **Distribution Prediction:** More sophisticated models estimate the full response distribution 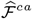. While these distributions typically lack closed-form expressions, one can draw samples 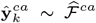 and estimate **x**^*ca*^ using 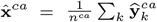.

### Limitations of Traditional Performance Metrics in Perturbation Response Prediction

When measuring the accuracy of prediction models, it is common to use a training–testing strategy. For the set of observed experiments 𝒪 = {**Y**^*ca*^ : (*c, a*) ∈ Ω}, we can partition 𝒪 into a training set 𝒯 and a testing or held-out set ℋ, such that

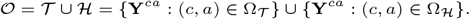

Models are trained on 𝒯 and the prediction results are evaluated on ℋ.

For continuous outcomes, several metrics are widely used to assess how closely predicted values match with the ground truth in the testing set. These metrics include mean absolute error (MAE), mean squared error (MSE), root mean squared error (RMSE), and *R*^2^. Generally, these metrics can be represented as a function 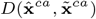, where they quantify the difference between in vitro mean 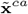 (in a held-out test set) and model predictions **x**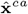, which are learned from the training data, Figure 1B. As an example, the *R*^2^ performance measure for a test experiment (*c, a*) ∈ ℋ can be written as:

**Fig. 1.**
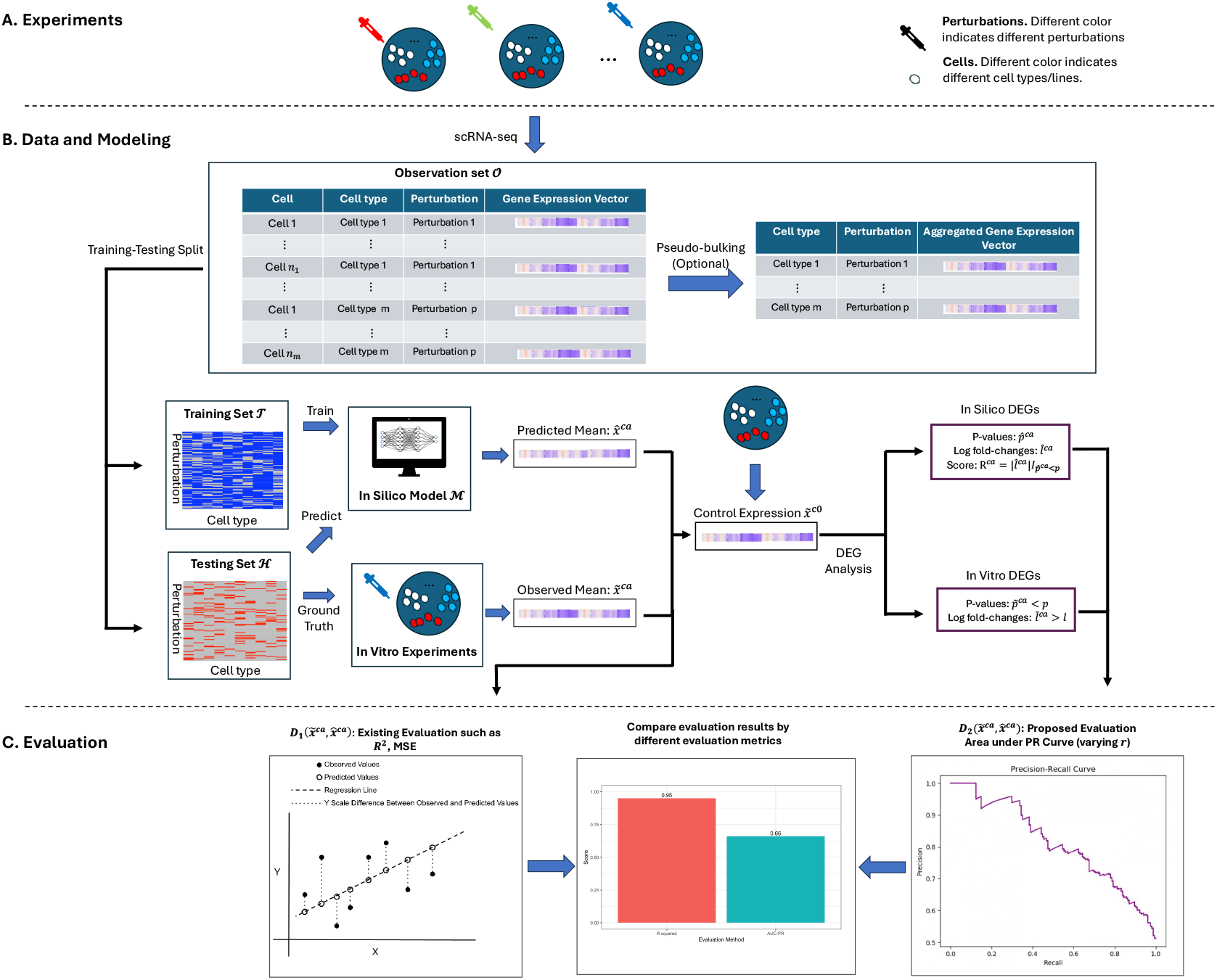
An illustration of the procedure. **A**. Cellular perturbation experiments are conducted, and responses are measured using single-cell RNA sequencing (scRNA-seq). **B**. The resulting single-cell expression data are then used to train in silico models. **C**. Different evaluation frameworks may yield different conclusions regarding model performance.

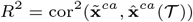

where 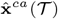 is the predicted expression vector for the held-out experiment (*c, a*), extrapolated from the training set 𝒯.

While *R*^2^ has become the de facto standard for evaluating gene expression prediction models [18, 17, 19, 32, 29, 5], it presents several limitations in the context of cellular perturbation experiments. These limitations stem from the unique characteristics of gene expression data and the biological nature of cellular responses to perturbations:

- **High Dimensionality and Sparsity:** Single cell gene expression data typically encompasses thousands of genes, with many showing little or no expression (zeros) in most conditions [32, 19]. A model can achieve high *R*^2^ values simply by correctly predicting these consistently non-expressed genes, without capturing biologically meaningful patterns.
- **Limited Perturbation Effects:** Perturbations typically affect only a small subset of genes directly, with the majority showing minimal expression changes [2]. As a global metric, *R*^2^ can be dominated by the large number of unaffected genes, potentially masking poor performance in predicting the crucial differentially expressed genes.
- **Biological Relevance:** A high *R* score may result from accurate predictions of baseline expression levels or housekeeping genes, while failing to capture perturbation-induced changes that are most relevant for biological interpretation and downstream analysis.

To address these limitations, researchers have proposed few complementary evaluation strategies. For instance, [19] evaluated *R*^2^ specifically on the top 100 differentially expressed genes, defined as genes showing significant expression changes between the experiment (*c, a*) and the unperturbed state (*c*, 0), while [29] computed *R*^2^ on the differential expression scale. As we demonstrate in this work, these auxiliary metrics still provide only indirect assessment of a model’s ability to predict biologically meaningful perturbation effects. A more comprehensive evaluation framework is needed to directly assess how well models capture the specific gene expression changes that are most relevant for biological interpretation and downstream analysis.

### Assessing Model Performance via Differential Expression Classification

Cellular perturbation datasets are primarily used for differential expression analysis, which is essential for identifying signature markers in response to various perturbations across specific cell types or cell lines. For example, the LINCS project [15] conducts such experiments and, after several processing steps, generates signatures representing differential expression in their level 5 processed data. Given this, a more effective approach to evaluate in silico model performance is to assess how accurately these models can replicate the differential expression outcomes derived from in vitro data. Building on this idea, we propose a new evaluation method that incorporates differential expression analysis using the in silico model prediction results to better assess the model’s ability to identify signature markers.

Standard differential expression analysis involves two key steps. The first step is conducting a hypothesis test between two groups of gene expression samples under different conditions to compare their mean expression differences. This test returns a p-value, indicating the statistical significance of the mean differences. The second step calculates the log fold-change by comparing the mean expressions between the two groups.

For in vitro experiment in context *c*, when comparing the expressions for gene *g* between experiment (*c, a*) and control condition (*c*, 0), the negative log_10_ p-value denoted by 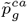 could be obtained by the following hypothesis test:

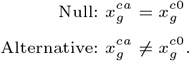

Here, 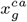 and 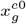 are the mean gene expressions under (*c, a*) and (*c*, 0), respectively, as defined in our notation. The log fold-change for gene *g* is defined by

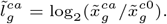

By applying appropriate cutoffs for the negative log_10_ p-value and absolute value of the log_2_ fold-change, denoted as *p* and *l*, DEGs can be identified as those surpassing both thresholds. In other words, with in vitro data, genes may be labeled as DE or non-DE using the indicator random variable 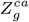 as follows:

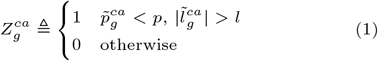

For in silico model predictions, calculating the log fold-change is straightforward and can be expressed as

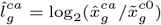

since we are comparing the predicted post-perturbation expression with the in vitro control expression. However, calculating p-values for in silico predictions presents unique challenges, as most models only predict point estimates without uncertainty quantification. For models that estimate distributions 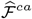, p-values 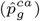 can be computed by comparing samples from the predicted distributions against observed control expressions.

To evaluate model performance in identifying DEGs, we propose a ranking-based approach that generates a family of classifiers with varying stringency. For each gene *j* under experiment (*c, a*), we define a ranking score that combines the magnitude of expression change with statistical significance:

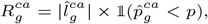

where 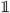 is the indicator function.

For any threshold *r* on this ranking score, we can define a classifier:

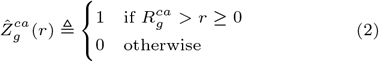

This formulation creates a continuous spectrum of classifiers, from stringent (high *r*, few predicted DEGs) to permissive (low *r*, many predicted DEGs). We evaluate these classifiers using precision-recall (PR) curves. The PR curve analysis is particularly suitable for differential expression analysis due to the inherent class imbalance, where DEGs typically constitute a small fraction of all genes. The precision-recall curve is generated by iterating through the gene list sorted according to their rank score, using each score as a threshold to calculate precision and recall at that level. This results in a set of points in 2-dimensional precision-recall space. The PR curve is then constructed from these points using a non-linear interpolation technique as described in [10]. The AUC-PR is the area under the PR curve, providing a quantitative summary of model performance in identifying DEGs.

We define the baseline AUC-PR as

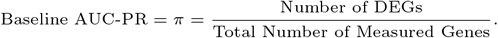

This baseline is achieved by two models which represent no predictive ability: First, a classifier which ranks all genes the same 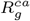 equal for all *g*), obtains an AUC-PR of *π*. Second, a classifier which produces random ranks, independent of the in vitro differential expression status 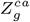, has an expected precision of *π* at every recall value and thus an expected AUC-PR of *π*. Thus the Baseline AUC-PR serves as a minimum performance threshold - any useful model must achieve an AUC-PR above this level.

Our complete evaluation procedure consists of three steps:

1. Establish ground truth 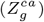 by identifying DEGs from in vitro data using standard thresholds based on equation (1),
2. Compute ranking scores 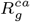 from in silico predictions and generate a family of classifiers 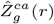 by varying threshold *r* using equation (2), **3)** Evaluate model performance using precision-recall analysis, comparing against both the ground truth labels and the baseline AUC-PR (*π*).

This framework provides a direct assessment of a model’s ability to identify biologically meaningful expression changes, addressing the limitations of traditional evaluation metrics like *R*^2^.

## Results

### DEG Prediction on Cell-level Responses Under Single Stimulus Across Multiple Cell Types

In this section, we consider the PBMC single cell perturbation dataset introduced in [14]. The study measured gene expression for human peripheral blood mononuclear cells (PBMCs) under two conditions: a control condition and a stimulated condition with interferon gamma. The dataset was further processed by [19], including gene filtering, normalization, and log-transformation. The final data includes 18,868 cells across 7 cell types, with 6998 highly variable genes measured for each cell as the cellular responses. Using the notations introduced in previous sections, let 𝒞 = {1, 2, …, 7} represent the cell types and 𝒜 = {0, 1} represents the conditions, where 0 denotes the control condition and 1 denotes the stimulated condition.

Previous work from [19] compared model performance between scGen and other modeling approaches, including linear benchmark models and some deep learning alternatives. These models are applied for out-of-sample prediction on the stimulated condition for each cell type. To evaluate model performance generally, the models are trained 7 times, with each iteration holding out cells measured in the stimulated condition of one of the seven cell types as the testing set. [19] assessed model performance using *R*^2^ between the real stimulated and the predicted mean gene expression for both all 6998 genes and the top 100 DEGs.

In our study, we evaluate the performance of different models in identifying differentially expressed genes (DEGs). We compare three approaches: (**1**) a single-factor linear regression model that includes only **cell type** as a categorical predictor (referred to as the cell type model), (**2**) a two-factor linear regression model that includes both **cell type** and **condition** as categorical predictors (referred to as the two-factor model), and (**3**) scGen, a deep generative model [19].

Differentially expressed genes are identified using a log_2_-fold change threshold of *l* = 0.3 and a p-value threshold of 10^*−*10^. Table 2 summarizes the number of DEGs detected in each cell type under these criteria. Additionally, it provides the number of cell samples per condition for each cell type and reports the baseline area under the precision-recall curve (the number of DEGs divided by 6998).

**Table 1.** Summary of recently developed in silico models and evaluation metrics used in testing.

| Year | Model | Metric |
| --- | --- | --- |
| 2019 | scGen [19] | $R^2$ |
| 2019 | XGBoost [16] | MAE |
| 2020 | trVAE [17] | $R^2$ |
| 2021 | Cellbox [33] | $R^2$ |
| 2021 | CPA [18] | $R^2$ |
| 2021 | ENformer [5] | $R^2$ |
| 2022 | SI-A [32] | $R^2$, RMSE |
| 2022 | Ensemble [4] | $R^2$ |
| 2022 | GEARS [29] | $R^2$, MSE |
| 2024 | scFoundation [13] | $R^2$, PCC |
| 2024 | scGPT [8] | $R^2$ |

**Table 2.** Overview of processed data from [19]

| Cell Type | Condition | # of cells | # of DEGs | Baseline AUC-PR |
| --- | --- | --- | --- | --- |
| CD4-T | Control | 2437 | 30 | 0.004 |
|  | Stimulated | 3127 |  |  |
| B | Control | 818 | 40 | 0.005 |
|  | Stimulated | 993 |  |  |
| CD8-T | Control | 574 | 30 | 0.004 |
|  | Stimulated | 541 |  |  |
| CD14+Mono | Control | 1946 | 101 | 0.014 |
|  | Stimulated | 615 |  |  |
| NK | Control | 517 | 31 | 0.004 |
|  | Stimulated | 646 |  |  |
| Dendritic | Control | 615 | 88 | 0.013 |
|  | Stimulated | 463 |  |  |
| FCGR3A+Mono | Control | 1100 | 98 | 0.014 |
|  | Stimulated | 2501 |  |  |

### High *R*^2^ Does Not Imply Ability to Identify Differentially Expressed Genes

Our analysis reveals a fundamental limitation of *R*^2^ as an evaluation metric for model performance in the context of differential expression analysis. This limitation becomes particularly evident when evaluating the single-factor *cell type model*, which achieves high *R*^2^ scores while failing to capture differential expression patterns.

#### Global *R*^2^ Performance Masks Poor DEG Prediction

Figure 2 demonstrates this limitation by comparing the *cell type model* ‘s performance using two perspectives: *R*^2^ across all genes versus *R*^2^ restricted to DEGs. Figure 2A shows the correlation between predicted and actual stimulated expressions for CD4 T cells, where the model achieves an *R*^2^ of 0.87. However, a closer examination reveals that predictions for non-DE genes (blue dots) show substantially higher correlation with the ground truth compared to predictions for DE genes (red dots).

**Fig. 2.**
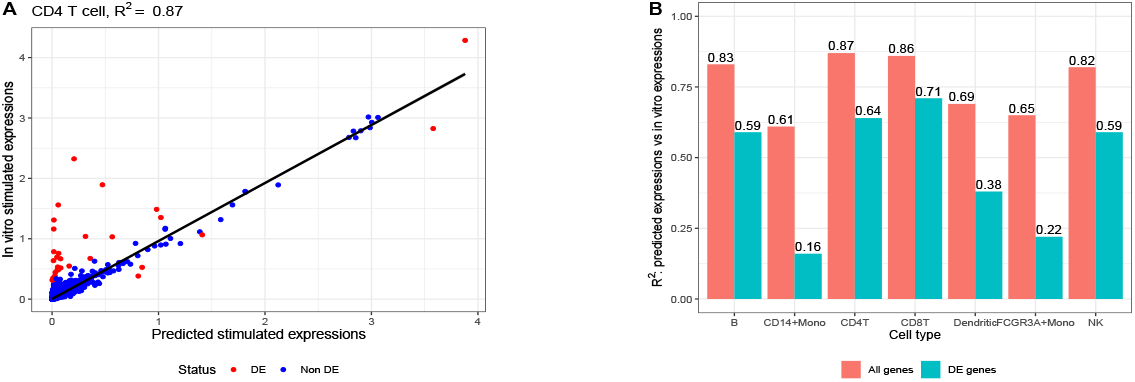
*R*^2^ **Performance of the *Cell Type Model*. A**. Scatter plot of predicted vs. actual stimulated expressions for CD4 T cells. The overall *R*^2^ across all genes is 0.87. *Note that in the cell type model, the predicted stimulated expression for each gene is simply the average expression in the unperturbed state*. **B**. Comparison of *R*^2^ computed across all genes and *R*^2^ computed across DEGs for all cell types.

Figure 2B quantifies this disparity further by comparing *R*^2^ computed over all genes versus DEGs across all cell types. The results show consistently higher *R*^2^ values when computed using all genes compared to DEGs, with substantial variation across cell types. This pattern indicates that the model’s high overall *R*^2^ primarily reflects its ability to predict non-DE genes rather than its capacity to capture true differential expression.

#### Analysis of the Cell Type Model Failure

The *cell type model* ‘s inability to identify DEGs can be explained by its fundamental architecture. As a simple linear regression model that considers only cell type information, its prediction for a perturbed condition in a given cell type, denoted as 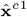 for experiment (*c*, 1), is given by:

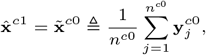

where 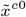 represents the mean expression of the target cell type under the control condition.

This formulation implies that the *cell type model* inherently assumes that average gene expression remains unchanged under perturbation. Consequently, differential expression analysis based on its predictions is meaningless: the predicted log fold-changes between true control and predicted perturbed conditions are uniformly zero, making differential expression analysis based on its predictions impossible, and the model produces no DEGs. Following our notation in the previous section, 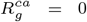 for all *g*, the only possible precision-recall value pair is (*π*, 1), achieved by labeling all genes as differentially expressed. The resulting AUC-PR is *π*, the baseline which indicates no predictive ability.

#### Implications for Model Evaluation in Differential Expression Analysis

This analysis highlights a critical limitation of using *R*^2^ as the sole evaluation metric for models intended for differential expression analysis. While *R*^2^ effectively captures global correlation between predicted and observed expressions, it provides limited insight into a model’s ability to identify differential expression patterns. A model can achieve an impressive *R*^2^ of 0.9 while completely failing to identify DEGs. The AUC-PR is also demonstrably more informative than *R*^2^ fit on DEGs. While *R*^2^ restricted to DEGs can be as high as 0.71 for CD8T cells, the AUC-PR is equal to *π* for all cell types, clearly indicating lack of any predictive ability of the model.

These findings emphasize the importance of using biologically relevant evaluation metrics, particularly when the downstream applications focus on DEG identification, such as biomarker discovery or experimental design. Metrics like AUC-PR, which explicitly evaluate a model’s ability to prioritize DE genes over non-DE genes, provide more meaningful assessment of performance in differential expression analysis tasks.

### Comparative Analysis of Two-Factor and scGen Models for DEG Prediction

We introduce a linear regression-based two-factor model and demonstrate its application to differential expression analysis. We also develop a statistical framework for generating p-values from scGen predictions, extending its capabilities beyond single-point predictions of perturbed gene expression. This enables a direct comparison between these fundamentally different approaches to DEG prediction.

#### The Two-Factor Model for DEG Detection

The two-factor model extends the cell type model by incorporating condition as a covariate in the linear regression framework. For a given cell type *c* under stimulated condition, we denote its prediction as 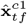, where subscript *tf* indicates the two-factor model estimate.

The two-factor model assumes that log-transformed expression data follows:

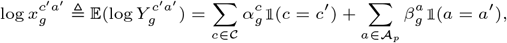

where 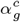 represents the baseline expression in cell type *c* and 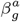 captures the effect of stimulation *a* for gene *g* with 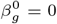 for control condition.

To identify DEGs, we test the hypothesis:

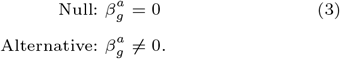

The predicted expressions for all genes are given by the vectors 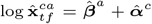 for perturbed and 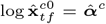 for control conditions. The log_2_ fold change vector is proportional to 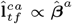, with rank score for each gene 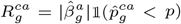, where 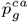 represents the negative log_10_ p-value from test (3).

#### scGen: A Deep Learning Approach to DEG Prediction

While scGen [19] does not inherently provide DEG predictions, we developed a systematic approach for DEG identification. As a variational autoencoder-based model, scGen learns the distribution of gene expression under stimulation, denoted as 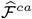 for cell type *c* under stimulation. We generate samples 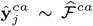 where 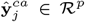 and 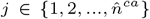.

The predicted mean expression is computed as:

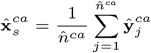

where subscript *s* denotes scGen estimates.

The log fold-change is calculated as 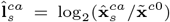 with statistical significance assessed via two-sample t-tests between 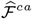 samples and control condition samples.

#### Comparative Performance Analysis

Our analysis of CD4T cell predictions reveals intriguing performance patterns. Figure 3A shows that scGen achieves a higher *R*^2^ (0.9) compared to the two-factor model (0.87). However, the two-factor model’s performance is remarkable given its simplicity compared to scGen’s sophisticated deep learning architecture, suggesting that high *R*^2^ may be achievable with relatively simple models.

The log fold-change analysis (Figure 3B) provides deeper insights, with dashed lines at *l* = 0.3 delineating the log_2_ fold change threshold used for differential expression boundaries. The two-factor model demonstrates superior performance in fold-change prediction (*R*^2^ = 0.64 vs. scGen’s 0.54). Figure 3C presents precision-recall curves for DEG classification, where the two-factor model achieves an AUC-PR of 0.62, outperforming scGen’s 0.55. While both substantially exceed the baseline AUC-PR (0.004), their moderate performance suggests significant room for improvement in DEG identification.

**Fig. 3.**
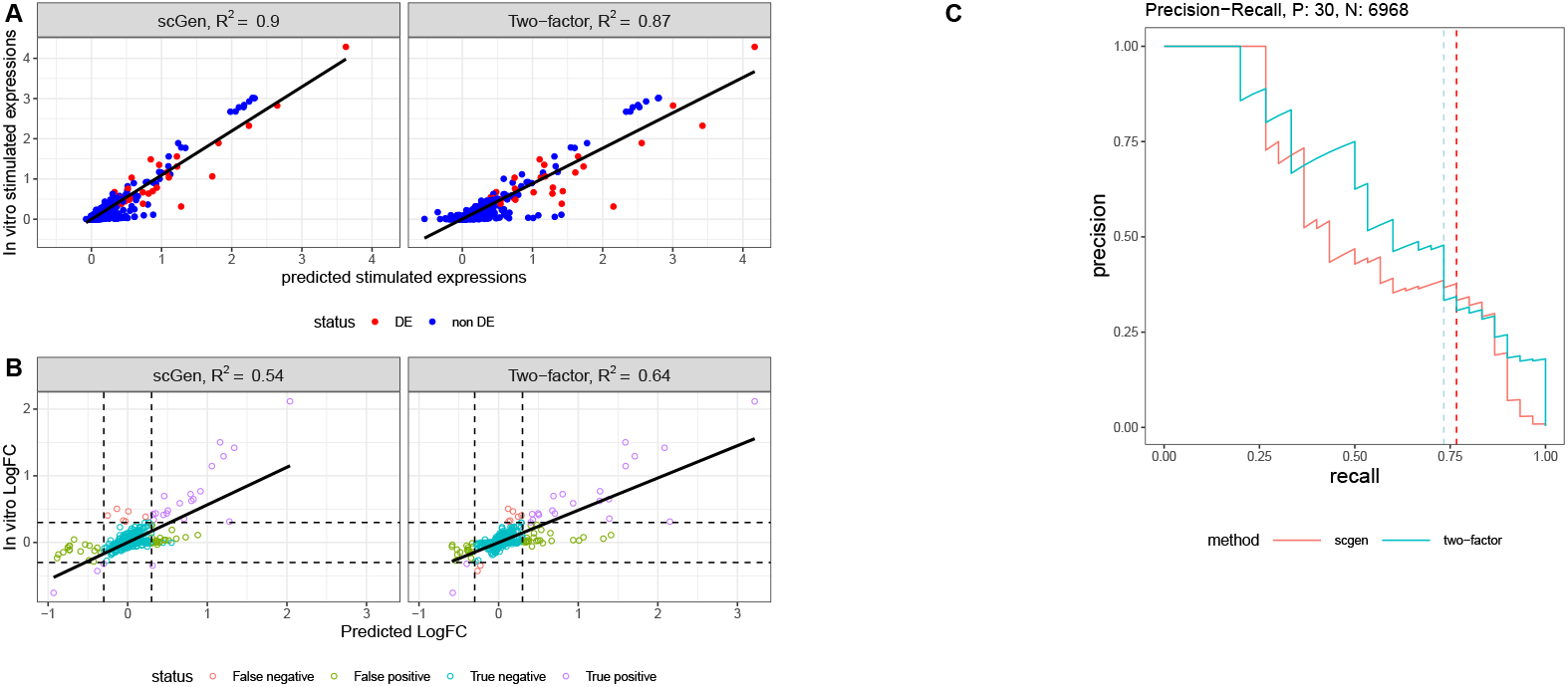
Performance analysis of CD4T stimulation predictions. **A**. Predicted versus actual stimulated expression scatter plot for CD4T cells. **B**. Predicted versus actual log fold-change comparison between control and stimulated conditions. **C**. Precision-recall curves for DEG prediction performance, with dashed lines indicating recall at 0.3 log fold-change threshold.

#### Cross-Cell Type Performance and Implications

The cross-cell type analysis (Figure 4) reveals consistent patterns. Figure 4A shows both models achieving high *R*^2^ values across cell types, but Figure 4B’s AUC-PR analysis exposes limitations in DEG identification capabilities.

**Fig. 4.**
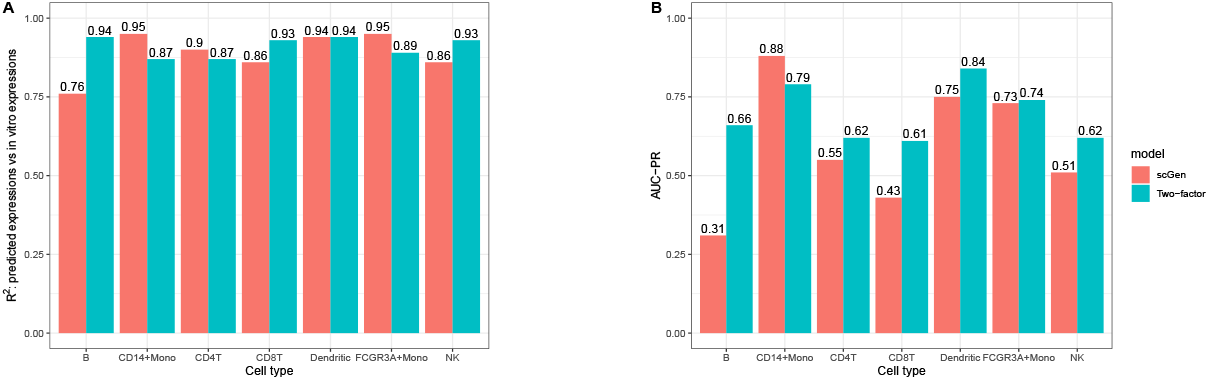
Cross-cell type performance comparison. **A**. *R*^2^ performance metrics across cell types. **B**. AUC-PR performance metrics across cell types.

Table 3’s precision analysis at various recall thresholds (25%, 50%, 75%) provides practical insights into model reliability. The similar performance between scGen and the two-factor model, coupled with moderate precision levels, suggests that current methods face significant challenges in reliable DEG identification. This finding has important implications for experimental design and validation strategies in differential expression studies.

**Table 3.** Precision values at specific recall levels for scGen and two-factor models in each cell type

| Cell Type | Method | Precision |  |  |
| --- | --- | --- | --- | --- |
|  |  | at 25% Recall | at 50% Recall | at 75% Recall |
| CD4-T | scGen | 1 | 0.44 | 0.38 |
|  | Two-factor | 0.89 | 0.75 | 0.34 |
| B | scGen | 0.41 | 0.39 | 0.18 |
|  | Two-factor | 1 | 0.67 | 0.49 |
| CD8-T | scGen | 0.73 | 0.37 | 0.22 |
|  | Two-factor | 0.8 | 0.65 | 0.5 |
| CD14+Mono | scGen | 0.93 | 0.91 | 0.88 |
|  | Two-factor | 1 | 0.82 | 0.66 |
| NK | scGen | 0.73 | 0.48 | 0.4 |
|  | Two-factor | 0.89 | 0.62 | 0.47 |
| Dendritic | scGen | 0.76 | 0.8 | 0.78 |
|  | Two-factor | 1 | 0.88 | 0.73 |
| FCGR3A+Mono | scGen | 0.93 | 0.83 | 0.67 |
|  | Two-factor | 1 | 0.82 | 0.57 |

### DEG Prediction on Population-level Responses Under Multiple Perturbations Across Multiple Cell Types

The previous section emphasized the modeling and evaluation approach for predicting single-cell level gene expressions. However, scRNA-seq data often contains significant noise at the individual cell level. To address this, pseudo-bulking is commonly used to aggregate gene expression across cell groups, reducing variability and highlighting population-level perturbation effects. In this section, we apply in silico models to pseudo-bulked data, allowing us to evaluate the models’ ability to accurately identify DEGs at the population level, offering a more interpretable assessment of performance.

We utilized a dataset from the Kaggle Single-Cell Perturbations Competition [7], derived from a novel single-cell perturbation analysis of PBMCs. This dataset features gene expression profiles following treatment with 144 compounds selected from the LINCS Connectivity Map[7], with measurements taken 24 hours post-treatment. PBMCs were collected from three healthy donors. For each donor, cells were plated onto two 96-well plates, resulting in six plates total. Each plate included:

- **Positive controls:** Two rows of wells treated with Dabrafenib and Belinostat.
- **Negative control:** One row of wells treated with DMSO.
- **Perturbations:** The remaining wells (144 unique perturbations in total), with one perturbation applied per well.

Each well contains 6 cell types, including T cells (regular, CD4 positive and CD8 positive), B cells, NK cells, and myeloid cells, with approximately 300–400 cells per cell type. The dataset includes gene expression profiles for 18,211 genes across six cell types, and yielding a theoretical total of 882 possible context (cell type) action (perturbation) pairs. However, some pairs are missing. For example, only 15 perturbations are observed in B cells and myeloid cells, and four pairs are absent in CD8 positive T cells. As a result, the dataset comprises a total of 614 observed pairs.

### Pseudo-bulking and Preprocessing

#### Pseudo-bulking is applied to this dataset for population-level analysis

Under the experimental setup, let 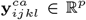 denote a single cell gene expression profile sample. Here, (*c, a*) represents the unique context (cell type) action (perturbation) pair, *i* ∈ {1, 2, 3} indicates the donor, *j* ∈ {1, 2, …, 6} indicates the plate, *k* ∈ {*A, B*, …, *H*} indicates the row (library) on a specific plate, while *l* represents the single cell sample. The subscripts {*ijk*} are collectively referred to as the **“location”** indicator, capturing batch effects associated with a (*c, a*) pair. Notably, these location indicators are not independent due to the experimental design. Each donor is assigned two specific plates: *j*|*i* = 1 ∈ {1, 2}, *j*|*i* = 2 ∈ {3, 4}, and *j*|*i* = 3 ∈ {5, 6}. Within each location defined by {*ijk*}, experiments are conducted across all cell types.

In this experiment, the first three columns (out of 12 total columns) on each plate are allocated for **control actions** (DMSO, dabrafenib, and belinostat). For a given control action and donor *i*, there are 16 possible locations for this control action across the two plates assigned to that donor. Each **perturbation action** is applied to one of the remaining wells on the plates assigned to the specific donor *i*. Consequently, for a given perturbation action *a*, once *i* is determined, the location {*ijk*} is uniquely specified. Under this setup, each action can have three possible locations, corresponding to the three donors *i* ∈ {1, 2, 3}.

We follow the data processing pipeline applied in the competition[7]. Pseudo-bulking is performed using sum aggregation for each cell type within a specific group defined by the location indicator {*ijk*}. Denote the pseudo-bulked expression profile as 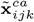. Then the pseudo-bulking procedure may be written as

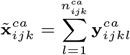

where 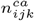 is the total number of single cell samples in group {*ijk*} for a given context-action pair (*c, a*). Ideally, pseudo-bulking results in 48 pseudo-bulked samples for each **control pair** (the context-action pair that formed with control actions) and 3 pseudo-bulked samples for each **perturbation pair** (the context-action pair that formed with perturbation actions). However, due to experimental constraints, some pairs contain fewer than three observations. During preprocessing, we excluded pairs with incomplete measurements across donors, resulting in a curated dataset where each remaining pair includes complete data from all three donors.

In vitro DE analysis was conducted using a linear model implemented with Limma [27], identical to the approach used in the competition [7]. This model aims to identify genes that are differentially expressed between specific perturbations and the negative control (DMSO) within each cell type. To adjust for batch effects, donor (*i*), plate (*j*), and library (*k*) are included as covariates in the model. Limma estimates ***α***^*c*^ and ***β***^*a*^ and their corresponding p-values.

### In silico models and Evaluation of performance

#### The dataset was divided into training and testing sets for evaluation of model performance

From pairs with complete responses across all donors, 100 were randomly selected for the testing set, while the remaining pairs formed the training set. On the training set, we applied three linear benchmark models and the SI-A model [32], independently for each donor on a log-normalized scale. The benchmark models included two single-factor linear regression models—**a cell type model** and **a perturbation model**—and **a two-factor** linear regression model. All the models applied on this datasets output a vector of predicted gene expressions 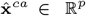 his approach generates predicted gene expression levels for each context–action pair within each donor. Importantly, within each donor, each perturbation is tied to a unique set of “location” parameters {i,j,k}. By assigning the predictions to the same “location” parameters as the ground truth, we preserve the experimental data structure. This consistency allows us to apply the same Limma pipeline, which includes donor (*i*), plate (*j*), and library (*k*) as covariates for in vitro DE analysis, to predicted expressions. Additionally, to derive model-predicted DEGs and their ranking scores, we first transformed the predictions back to the original count scale. We then applied the same Limma pipeline used in the in vitro differential expression analysis to compute p-values and log fold-changes.

Model performance is then evaluated across pairs in the testing set using multiple metrics. First, we calculate the *R*^2^ values between the observed and predicted gene expression levels across all 18,211 genes **for each donor**. The *R*^2^ values across donors provide an overall measure of the correlation between the predictions and the ground truth for each testing pair. To evaluate the model’s ability to identify DEGs, we compare the in silico DEG results to the in vitro DEG results, which serve as the ground truth. For each testing pair, we compute the PR curve and AUC-PR to quantify the model’s ability to accurately identify DEGs.

#### Comparative Performance Analysis on Specific Testing Pair

We first evaluate model performance on a specific cell type–perturbation pair: T cells CD4 positive perturbed by Perhexiline. Figure 5 compares two evaluation metrics: *R*^2^ and the proposed PR-curve approach. Figure 5A presents a scatter plot comparing predicted gene expression (log scale) with in vitro gene expression (log scale) under donor 1. Notably, the *R*^2^ values for other donors are consistent with the *R*^2^ obtained for donor 1. The cell type model, two-factor model, and SI-A model all achieved high *R*^2^ values around 0.95, while the perturbation model had a lower *R*^2^ of 0.82. In the scatter plot, red points represent DEGs identified via in vitro DEG analysis, while blue points represent non-DE genes. Notice that, the correlation of predicted log expression and in vitro log expression among DEGs are lower than the overall correlation. The *R*^2^ values, complemented by the scatter plots, suggest that the cell type model, two-factor model and SI-A perform well in predicting overall gene expression levels in this specific pair.

**Fig. 5.**
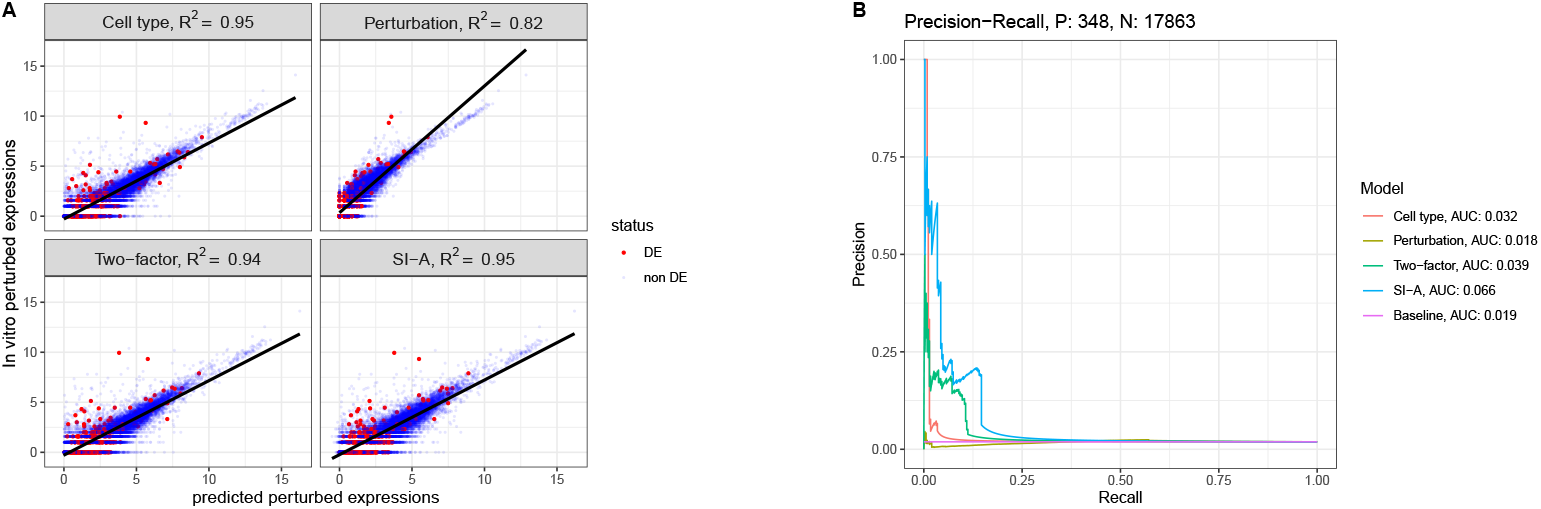
Predictions for T cell CD4+ perturbed by Perhexiline. **A**. Scatter plots of In vitro expression versus In Silico predictions for donor 1. **B**. PR-curves obtained by the four models compared with the baseline PR curve. Cell type model, two-factor model and SI-A outperform the baseline with slightly higher AUC-PR, while the perturbation model fails to outperform the baseline.

Figure 5B compares the PR-curves and the corresponding AUC-PR values for each model against the baseline performance, which represents random guessing. The cell type model, two-factor model, and SI-A all outperform the baseline, with SI-A achieving the highest AUC-PR of 0.066. This result indicates that, when evaluating the models’ ability to identify DEGs, as reflected by the AUC-PR metric, the performance is notably weaker. This discrepancy underscores the limitations of relying solely on *R*^2^ as an evaluation metric.

#### Comparative Performance Analysis across All Testing pairs

Figure 6 shows the evaluation of model performance across all 100 testing sets Figure 6A displays a dot plot illustrating the distribution of *R*^2^ values across all testing sets for each donor. The average *R*^2^ values follow a pattern similar to that observed in Figure 5C, with cell type model, two-factor model and SI-A achieving average *R*^2^ values close to 0.9, indicating consistent performance in predicting overall gene expression levels.

**Fig. 6.**
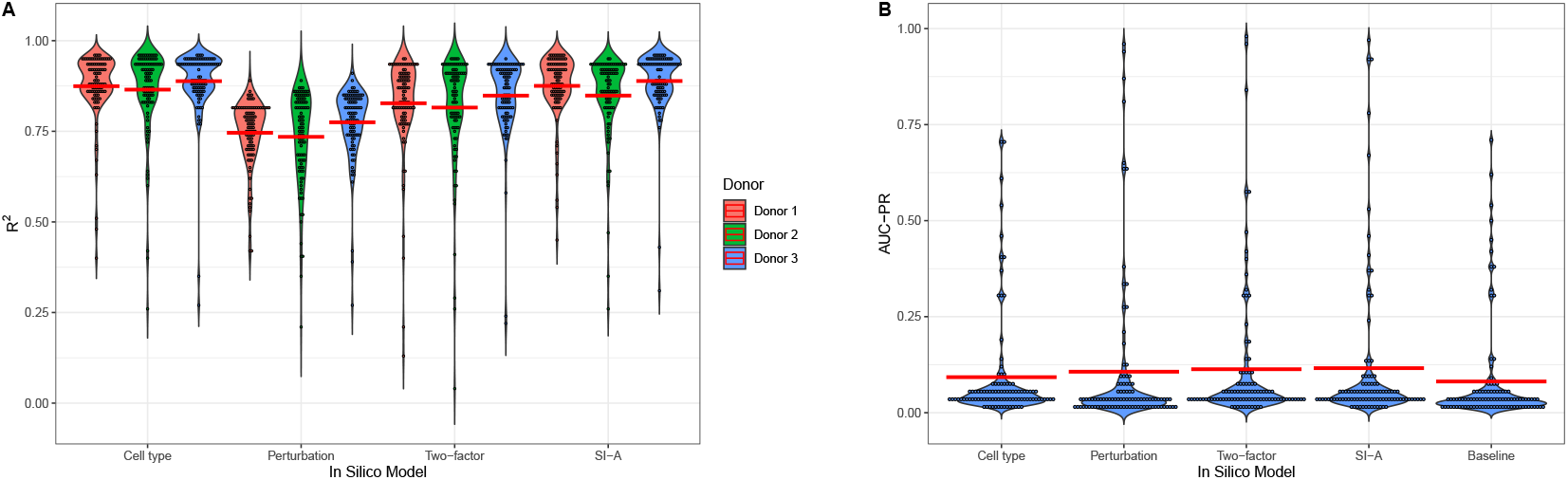
Predictions on 100 test pairs. **A**. Distribution of *R*^2^ values across all 100 testing sets for each donor. The red lines indicate average *R*^2^ for the models. **B**. Distribution of AUC-PR values for each model and baseline AUC-PR across all 100 testing sets. The red lines indicates average AUC-PR.

Figure 6B provides a dot plot showing the distribution of AUC-PR values across all testing sets. In contrast to the *R*^2^ metric, the average AUC-PR values for all models are considerably lower, clustering around 0.1. Among the models, SI-A slightly outperforms the others, achieving an average AUC-PR of 0.12. Interestingly, in some testing sets, the in silico models achieve AUC-PR values approaching 1. This exceptional performance in certain cases is primarily attributable to the presence of a large number of positive cases (DEGs) in the corresponding cell type–perturbation pairs, which can make DEG identification easier. These cases usually have high baselines as shown in the Figure 6.

#### Implications from Comparative Performance Analysis

These comparative results reinforce our earlier finding that *R*^2^, while effective at assessing overall prediction accuracy, does not account for a model’s ability to identify DEGs—a critical requirement for many biological applications for cellular perturbation experiments. Specifically, *R*^2^ evaluates the agreement between predicted and observed gene expression levels across all genes, but it lacks sensitivity to the complex nature of DEG identification tasks, where those DEGs were determined by log fold-changes and statistical significance thresholds. This insensitivity limits the utility of *R*^2^ for evaluation models aimed at identifying biologically significant patterns.

In contrast, the proposed AUC-PR metric directly addresses these challenges by quantifying the precision and recall of DEG predictions, offering a focused evaluation of a model’s performance in this critical task. By focusing on the ability to accurately identify DEGs, the AUC-PR metric provides a complementary perspective that highlights aspects of biological interpretation that *R*^2^ does not capture.

## Discussion

In this paper, we present a novel framework for evaluation in silico models in the context of cellular perturbation experiments, with a particular focus on their ability to identify DEGs. By comparing traditional metrics like *R*^2^ with our proposed AUC-PR metric, we showed the limitations of existing evaluation methods. Our findings reveal that *R*^2^, despite its widespread use for assessing overall prediction accuracy, is of limited value in assessing a model’s ability to identify DEGs. High *R*^2^ values observed for certain in silico models indicate strong correlation between predicted and observed gene expression levels across all genes. However, these models often exhibit low AUC-PR values, reflecting poor performance in identifying DEGs. This discrepancy underscores the necessity of incorporating complementary evaluation metrics to provide a more comprehensive and biologically relevant assessment of model performance.

Our work also provides a more detailed evaluation of the absolute performance of the models by analyzing precision at specific recall levels. Precision at a given recall level evaluates the proportion of correctly identified DEGs among all predicted DEGs when the model achieves a certain recall. Our analysis revealed that even at moderate recall levels, the tested models demonstrate limited precision, suggesting that in silico models are not ready to replace in vitro experimentation. For example, as illustrated in Figures 5 and 6, while SI-A slightly outperforms simple linear models, the improvement is minimal. This finding aligns with previous studies showing that many sophisticated models fail to significantly surpass simpler linear models in similar tasks [3]. Our results complement these studies, and our emphasis on identifying DEGs and introducing biologically informed evaluation metrics offers a novel perspective.

The findings of this study also have implications for the development and application of in silico models in cellular perturbation research. Accurate identification of DEGs is crucial for a wide range of downstream analyses[15], such as uncovering disease-specific biomarkers, understanding gene regulatory networks, and elucidating cellular mechanisms under perturbation[23]. These tasks are fundamental for advancing drug discovery, designing targeted therapies, and improving our understanding of complex biological systems[24]. Our results suggest that relying solely on traditional metrics like *R*^2^ may overestimate a model’s utility for these critical tasks. Incorporating biologically relevant evaluation metrics like our proposed AUC-PR into the assessment pipeline ensures a more interpretable understanding of model performance.

This study also highlights several avenues for future research to further enhance the utility and interpretability of in silico models in cellular perturbation experiments. First, developing a more theoretically grounded approach for DEG analysis using in silico model predictions is essential. This includes improving methodologies for calculating statistical significance (p-values), which currently rely heavily on assumptions that may not adequately account for the inherent uncertainty in model predictions. Future work should focus on quantifying prediction uncertainties into the DEG analysis framework, thereby improving the robustness and reliability of p-value calculations.

Additionally, exploring how specific data characteristics, such as the proportion of DEGs, noise levels, or dataset sparsity, influence model evaluation metrics can provide deeper insights into model performance and limitations. Understanding these relationships can help optimize data preprocessing and model training strategies, ultimately leading to better predictions and more accurate identification of DEGs. Such efforts could also guide the design of more tailored evaluation protocols, ensuring that models are assessed under conditions that closely reflect real-world biological scenarios.

Finally, extending the proposed evaluation framework to include a broader range of datasets and prediction models will be critical for generalizing our findings. Incorporating diverse datasets with varying experimental conditions, cell types, and perturbation types will allow for a more comprehensive assessment of model performance. Additionally, benchmarking emerging in silico models, such as the transformer based scGPT[8], within this framework will help clarify its potential value in biological research. These efforts will not only validate the proposed evaluation framework but also guide the development of more robust and biologically meaningful predictive models for cellular perturbation research.

## Competing interests

No competing interest is declared.

## Data and Code Availability

The data and code could be found at https://github.com/hxzhu491/Cell-Perturbation-evaluation-Metric.

## Acknowledgments

NIH [R00HG011367 to A.A. and L.A.]. JPL received support from the National Cancer Institute and the National Center for Advancing Translational Sciences of the NIH [CCSG P30CA016672-46 and CCTS UM1TR004906].

## Notes

https://github.com/hxzhu491/Cell-Perturbation-evaluation-Metric

### Competing Interest Statement

The authors have declared no competing interest.

